# One pathway, two cyclic pentapeptides: heterologous expression of BE-18257 A-C and pentaminomycins A-E from *Streptomyces cacaoi* CA-170360

**DOI:** 10.1101/2020.10.23.352575

**Authors:** Fernando Román-Hurtado, Marina Sánchez-Hidalgo, Jesús Martín, Francisco Javier Ortiz-López, Daniel Carretero-Molina, Fernando Reyes, Olga Genilloud

## Abstract

The strain *Streptomyces cacaoi* CA-170360 produces the cyclic pentapeptides pentaminomycins A-E and BE-18257 A-C, two families of cyclopeptides synthesized by two nonribosomal peptide synthetases encoded in tandem within the same biosynthetic gene cluster. In this work, we have cloned and confirmed the heterologous expression of this biosynthetic gene cluster, demonstrating that each of the nonribosomal peptide synthetases present in the cluster is involved in the biosynthesis of each group of cyclopeptides. In addition, we discuss the involvement of a stand-alone enzyme belonging to the Penicillin Binding Protein family in the release and macrocyclization of the peptides.

## 2. Introduction

Cyclic peptides are one of the most important chemical classes of biomolecules with potential therapeutic applications. The cyclic polypeptide chain is formed by amide bonds between proteinogenic or nonproteinogenic amino acids, with a structure that confers reduced conformational flexibility, resistance to exo- and endopeptidases, increased cell permeability and better biological activities compared with their linear counterparts. Consequently, these molecules show in general low toxicity, good binding affinity and target selectivity (Abdalla & McGaw, 2018; Jing & Jin, 2020).

Bacterial cyclic peptides exhibit a wide variety of biological activities. Best examples of largely used cyclic peptides as therapeutic agents are the antibiotics gramicidin, vancomycin, daptomycin, and polymyxin B (Mogi & Kita, 2009; Levine, 2006; Tedesco & Rybak, 2004). In the last years, new bacterial bioactive cyclopeptides have been reported, such as pargamicins A-D (Igarashi et al., 2008; Hashizume et al., 2017), cyclotetrapeptides cyclo-(Leu-Pro-Ile-Pro) and cyclo(Tyr-Pro-Phe-Gly) (Abdalla, 2016), the cyclic pentapeptides BE-18257BE-18257 A-D (Miyata et al., 1992a; 1992b) and the pentaminomycins A-E (Jang et al., 2018; Kaweewan et al., 2020; Hwang et al., 2020). Cyclic pentapeptides BE-18257 A-D, were first isolated from *Streptomyces* sp. 7338 as endothelin receptor antagonists (Miyata et al., 1992a; 1992b). The pentaminomycins are cyclic pentapeptides that possess the common core sequence Val-Trp-N^δ^-OHArg and show distinct biological activities. Pentaminomycin A has antimelanogenic activity against alpha-melanocyte stimulating hormone (α-MSH)-stimulated B16F10 melanoma cells (Jang et al., 2018) and pentaminomycin C is active against Gram positive bacteria but no against Gram negative bacteria (Kaweewan et al., 2020). Both pentaminomycins C and D act as autophagy inducers on HEK293 cells (Hwang et al., 2020).

Recently, Kaweewan et al. (2020) and Hwang et al. (2020) proposed a biosynthetic gene cluster (BGC) for the production of pentaminomycins C-E and the cyclic pentapeptides BE-18257 A-B. The proposed cluster harbors regulatory, transport-related and biosynthetic genes, including cytochromes P450 and two NRPS, each of them predicted to synthesize one type of pentapeptide. The cluster lacks any gene harboring a thioesterase (TE) domain but contains a stand-alone enzyme belonging to the Penicillin Binding Protein (PBP) family, that, likewise to what has been demonstrated for SurE in the biosynthesis of surugamide (Matsuda et al., 2019a; 2019b; Zhou et al., 2019), is proposed to act as a trans-acting TE cyclizing both BE-18257 A-B and pentaminomycins C-E (Kaweewan et al., 2020; Hwang et al., 2020).

The strain *Streptomyces cacaoi* CA-170360 from MEDINA’s microbial collection produces the cyclic pentapeptides BE-18257 A-C (Figure 1) and pentaminomycins A-E (Figure 2) (Carretero-Molina et al., unpublished results). This strain has also been recently described as the producer of cacaoidin, the founding member of class V lanthipeptides (lanthidins) (Ortiz-López et al., 2020; Román-Hurtado et al., 2020). In this work, we demonstrate that pentaminomycins A-E and BE-18257 A-C are synthesized in this strain by a pathway highly similar to that described recently in other *S. cacaoi* strains (Kaweewan et al., 2020; Hwang et al., 2020). To that end, we have cloned and heterologously expressed both the complete BGC and a partial pathway that lacks the NRPS encoding pentaminomycins. In the last case, only BE-18257 antibiotics are detected, confirming the involvement of each NRPS in the biosynthesis of the respective cyclopentapeptides and the putative involvement of the pathway-located PBP-TE in the cyclization of both types of compounds.

**Figure 1.**
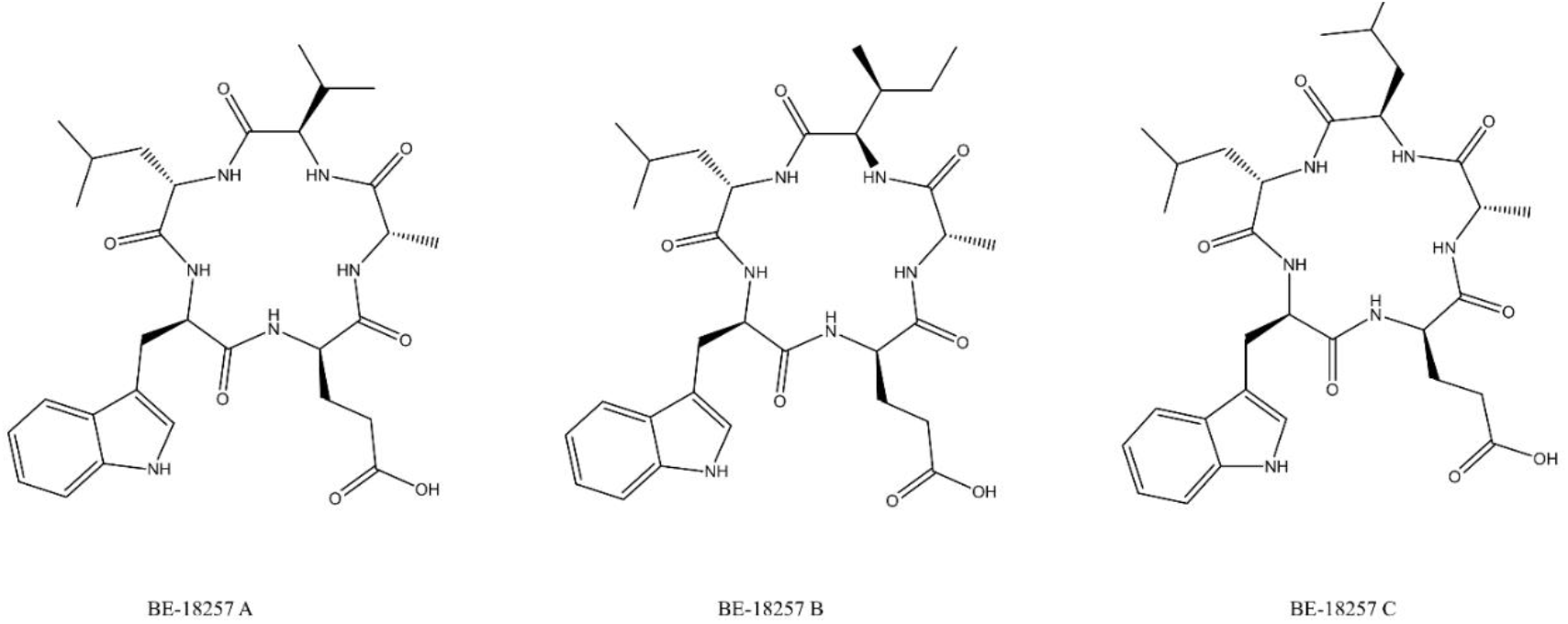
Structures of BE-18257 A-C antibiotics.

**Figure 2.**
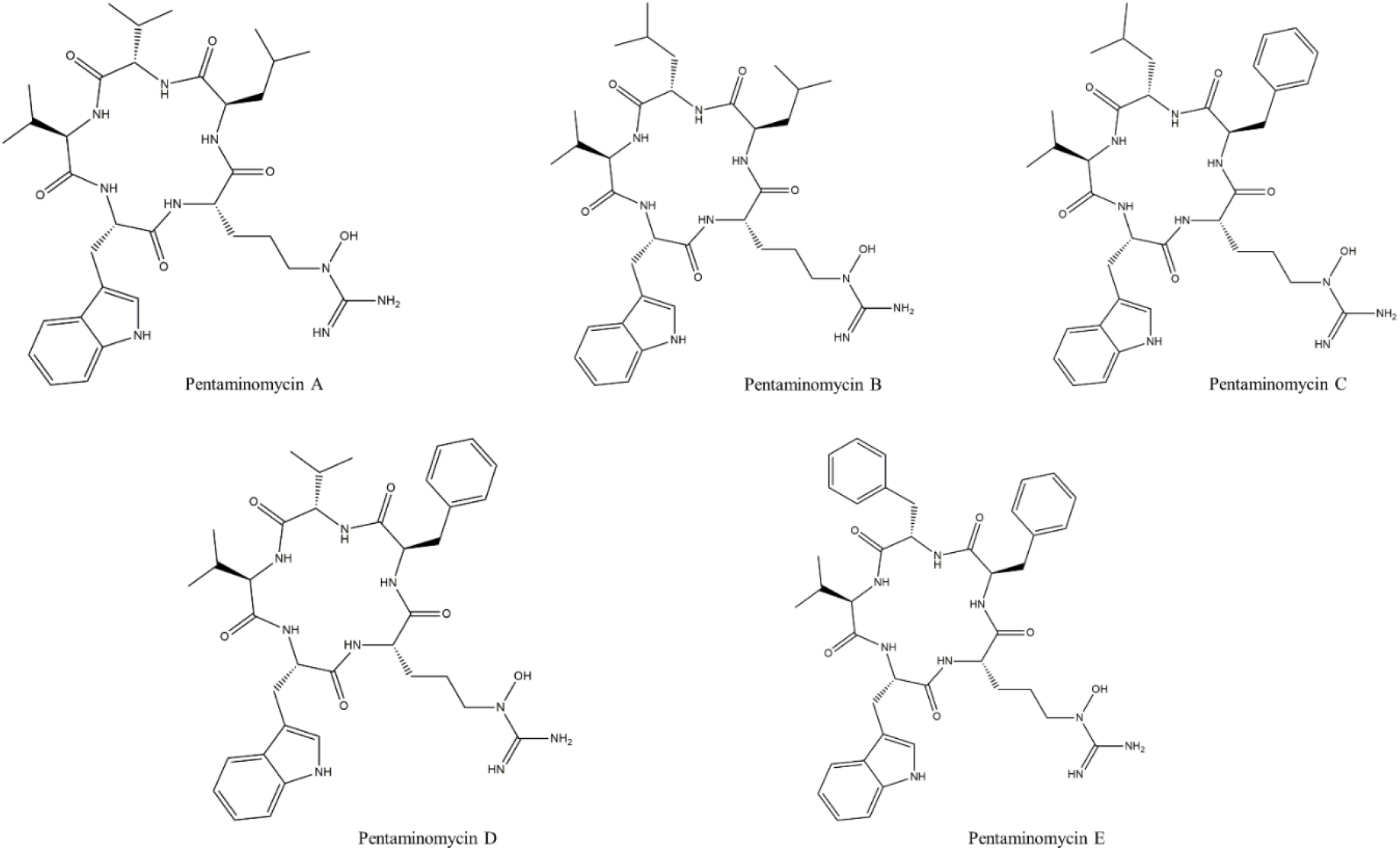
Structures of pentaminomycins A-E

## 3. Materials and methods

### 3.1. Bacterial strains and plasmids

Strain *Streptomyces cacaoi* CA-170360 from Fundación MEDINA’s culture collection was isolated from the rhizosphere of a specimen of *Brownanthus corallinus*, in the region of Namaqualand (South Africa). Electrocompetent NEB 10-β *E. coli* (New England BioLabs, Ipswich, MA, USA), *E. coli* ET12567 (LGC Standards, Manchester, NH, USA) and *E. coli* ET12567/pUB307 (generously provided by J.A. Salas) were employed throughout plasmids transformation and conjugation processes. Vector pCAP01 was used for the cloning of the BGCs. This plasmid is a *S. cerevisiae*/*E. coli*/actinobacteria shuttle, kanamycin resistant vector with a site-specific φC31 integrase which allows the incorporation of the cloned cluster to the genome of heterologous hosts (Zhang et al., 2019). *Streptomyces albus* J1074 was used as heterologous host (Chater & Wilde, 1980) and was kindly provided by J. A. Salas.

### 3.2. Growth and culture conditions

Culture media composition is described in the Supporting Information. *Streptomyces cacaoi* CA-170360 was cultured in ATCC-2 liquid medium and grown on an orbital shaker at 28°C, 220 rpm and 70% relative humidity. For the OSMAC approach, CA-170360 was cultured in six different liquid media (YEME, R2YE, KH4, MPG, FR23, DEF-15) and at three different times (7, 14 and 21 days). *E. coli* strains were routinely cultured in LB broth Miller (Sigma) (37 °C, 250 rpm), Difco LB agar Lennox (37 °C, static). Intergeneric conjugations were carried out on solid MA medium and. exconjugants were grown on antibiotic-supplemented MA plates. Antibiotics were added when required for selection of transformants at the following final concentrations: kanamycin (50 μg/mL), nalidixic acid (25 μg/mL), choramphenicol (25 μg/mL). For heterologous expression, MPG and R2YE media were used and the recombinant strains were incubated for 14 days on an orbital shaker at 28 °C, 220 rpm and 70% relative humidity.

### 3.3. Identification of *cpp* cluster from strain CA-170360 whole genome sequence

The genome sequence of *Streptomyces cacaoi* CA-170360 (Román-Hurtado et al, 2020) was analyzed by antiSMASH 5.1.2 (Blin et al, 2019) in order to find the biosynthetic gene cluster responsible for the production of pentaminomycins A-E and BE-18257 A-C. The *cpp* BGC sequence is available in the National Center for Biotechnology Information (NCBI) database under accession GenBank number MW038823.

### 3.4. Cloning and heterologous expression of the *cpp* gene cluster

The *cpp* cluster was cloned by CATCH (Cas9-Assisted Targeting of CHromosome), where a Cas9 endonuclease cleaves a large BGC guided by RNA templates (Jiang et al, 2015). Two clonings were performed in this work: one including the NRPS responsible to produce BE-18257 A-C and another one with both NRPS involved in the production of BE-18257 A-C and pentaminomycins A-E.

CRISPy-web tool (http://crispy.secondarymetabolites.org/) was employed to design 20 nt target sequences close to a PAM (Protospacer-Adjacent Motif) sequence ‘NGG’ (Tong et al, 2018) that is the target where Cas9 endonuclease cuts. Based on these sequences, the necessary primers are listed in Supporting Table 1. An overlapping PCR was carried out using three oligos, one target-specific oligo (Penta1-sgRNA, Penta2-sgRNA or Penta3-sgRNA) containing the target sequence and a T7 promoter and two universal oligos (sgRNA-F and sgRNA-R) in order to get the three Penta-sgRNAs needed for this study. Q5 High-Fidelity polymerase from New England BioLabs (Ipswich, MA, USA) was employed for this PCR. HiScribe T7 Quick Yield RNA synthesis kit (New England Biolabs) was used for the *in vitro* transcription and the products were purified by phenol/chloroform extraction and isopropanol precipitation, as Jian and Zhu described in their protocol (Jiang & Zhu, 2016).

*Streptomyces cacaoi* CA-170360 was cultured in ATCC-2 at 28°C, 220 rpm and 70% relative humidity to later in be embedded in low-melting agarose plugs where the in-gel Cas9 digestion was performed. The genomic DNA of the strain was extracted within the plugs using lysozyme, proteinase K and washing buffers, and once isolated the genome, in-gel digestion with Cas9 nuclease from *S. pyogenes* (New England BioLabs) was performed taking two plugs of agarose, a cleavage buffer (100 mM HEPES pH 7.5, 750 mM KCl, 0.5 mM EDTA pH 8, 50 mM MgCl_2_, DEPC-treated water) and the sgRNAs, and incubating at 37°C for 2 h. After the digestion, the agarose plugs were melted with a GELase treatment and the already digested DNA was recovered with an ethanol precipitation. The pCAP01 vector was previously amplified with the oligos pCAP01-Penta1-F/pCAP01-Penta1-R and pCAP01-Penta2-F/pCAP01-Penta2-R (Supporting Table 1) to get 30 nt overlapping ends. Then, the Cas9-cleavaged BGCs were cloned in the corresponding amplified vector by Gibson Assembly using a 2x Gibson Assembly Master Mix (New England BioLabs) and incubating at 50 °C for 1 h. The Gibson products, pCPP1 and pCPP2, were transformed into electrocompetent NEB-10-β *E. coli* cells. Plasmids pCPP1 and pCPP2 from isolated colonies were validated by restriction digestion with HindIII and NdeI.

As pCPP1 and pCPP2 contain the kanamycin-resistant marker, two triparental intergeneric conjugations were made using *E. coli* NEB-10/pCPP1 or *E. coli* NEB-10/pCPP2 and non-methylating Cm^R^ Km^R^ *E. coli* ET12567/pUB307 as donor strains, and spores of *S. albus* J1074 as recipient strain. For the negative control, *E. coli* NEB-10/pCAP01 and *E. coli* ET12567/pUB307 were used as donor strains.

Five positive transconjugants from each conjugation, together with the negative control and the wild-type strain *S. cacaoi* CA-170360, were grown on liquid MPG and R2YE media during 14 days at 28°C, and then acetone extracts from the cultures were obtained.

### 3.5. Extraction and detection of BE-18257 A-C and pentaminomycins A-E

Cultures of the recombinant strains *S. albus* J1074/pCPP1 and *S. albus* J1074/pCPP2, together with the negative control harboring empty pCAP01 vector and the original *S. cacaoi* CA-170360 as positive control, were subjected to extraction by liquid-liquid partition with acetone 1:1, stirring at 220 rpm for 2 hours. Once dried under a nitrogen atmosphere, the residue was resuspended in 20% DMSO/water and the resulting microbial extracts analyzed by LC-HRESI-TOF.

## 4. Results and Discussion

### 4.1. Production of cyclic pentapeptides by strain CA-170360

In our continuous effort to search for novel compounds, the strain *Streptomyces cacaoi* CA-170360 was shown to produce the cyclic pentapeptides BE-18257 A-C and, to a much lesser extent, the recently described pentaminomycins A-E after liquid fermentation in MPG medium for 13 days (Carretero-Molina et al., unpublished results). We followed an OSMAC approach (Bode et al, 2002) to identify the best production conditions of both families of cyclopeptides. The analysis included a total of six production media (YEME, R2YE, KM4, MPG, FR23 and DEF-15) and three fermentation times (7, 14 and 21 days). Production of BE-18257 A-C was the highest in KM4, MPG and FR23 media whereas the detection levels of pentaminomycins were very low. Pentaminomycins A-E were mostly produced in YEME and R2YE and required long incubations of 14 and21 days, although the production was still very low. Interestingly, in these conditions, BE-18257 A-C were produced in very small amounts, suggesting that these media ensure the biosynthesis of pentaminomycins to the detriment of the BE-18257 molecules (Supporting Figure 1). To our knowledge, CA-170360 is the first strain reported to produce all the five pentaminomycins described to date and the three BE-18257 compounds. The strain *Streptomyces* sp. RK88-1441 was shown to produce pentaminomycins A and B but no BE-18257 antibiotics (Jang et al, 2018) while only pentaminomycin C and BE-18257 A were isolated from the strain *Streptomyces cacaoi* subsp. *cacaoi* NBRC 12748^T^ (Kaweewan et al, 2020). Finally, Hwang et al, (2020) reported the production of pentaminomycins C-E and BE-18257 A-B from *Streptomyces* sp. GG23. These strains may have the capacity to synthesize all the pentaminomycins and BE-18257 antibiotic variants detected in strain CA-170360, but most probably the culture conditions used did not ensure the production of all the analogs.

### 4.2. Identification of the *cpp* gene cluster from the whole genome sequence of strain CA-170360

The whole genome sequence of *Streptomyces cacaoi* CA-170360 (Román-Hurtado et al, 2020) was analyzed with antiSMASH (Blin et al, 2019) and 31 putative BGCs, including NRPS, PKS and RiPPs, among others, were predicted (data not shown). One of the BGCs from contig 1 (*cpp* cluster) was identified as the putative pathway for the synthesis of both BE-18257 A-C and pentaminomycins A-E (GenBank number MW038823).

The *cpp* gene cluster contains 15 ORFs coding for proteins with the proposed functions shown in Table 1 and Figure 3. The gene organization and amino acid incorporation is highly similar to those previously proposed for these BGCs in strains NBRC 12748^T^ and GG23 (Kaweewan et al, 2020; Hwang et al, 2020), with two NRPS genes, each containing five adenylation (A) domains. The first NRPS gene (*cppB*) contains three epimerization (E) domains and a sequence of amino acids corresponding to Leu (A1), Trp (A2), Leu/Ser (A3), Ala (A4) and Val/Leu (A5). The second, third and fifth modules contain an epimerization domain, and they would be involved in the isomerization of an L- to D- amino acid, resulting in the final sequence L-Leu, D-Trp, D-Leu/Ser, L-Ala, D-Val/Leu, which is in accordance with the amino acid sequence of BE-18257 A-C (L-Leu,D-Trp,D-Glu,L-Ala,D-R) (Figure 4). Therefore, these results suggest that the first NRPS gene (*cppB)* may be involved in the biosynthesis of BE-18257 A-C antibiotics. Then, cyclization would complete the biosynthesis of the molecules. On the other hand, the second NRPS gene (*cppM*) contains two E domains and the sequence of amino acids incorporated would be Val/Leu/Phe (A1), Val (A2), Trp (A3), Arg (A4) and Leu/Phe (A5). As the second and fifth modules contain an epimerization domain, the final amino acid sequence would be L-Val/Leu/Phe, D-Val, L-Trp, L-Arg, D-Leu/Phe, which agrees with the amino acid sequence of pentaminomycins A-E (L-R1, D-Val, LTrp, L-N5-OH-Arg, D-R2) (Figure 5). Subsequent modifications such as hydroxylation and cyclization would complete the biosynthesis of the pentaminomycins. However, the *cpp* cluster also lacks a TE domain to release and cyclize the pentapeptides but contains a PBP-like stand-alone protein (*cpp*A) that may be involved in the release and cyclization of the peptide chains of both BE-18257 antibiotics and pentaminomycins, as it was proposed by Kaweevan (2020) and Hwang (2020). In fact, it has been recently described that SurE, a stand-alone enzyme belonging to the PBP family, is involved in the release and macrocyclization of two different surugamides (B and F) encoded in a single gene cluster (Matsuda et al, 2019a; 2019b; Zhou et al, 2019). This PBP-TE has been also reported in other NRP such as desotamide (Fazal et al, 2020), ulleungmycin (Son et al, 2017), noursamycin (Mudalungu et al, 2019), curacomycin (Kaweewan et al, 2017) or mannopeptimycin (Magarvey et al, 2006). The *cpp* cluster includes two ORFs (*cppI* and *cppJ*) encoding cytochrome P450 enzymes, which have been suggested to be involved in the N- hydroxylation of arginine to form 5-OH-Arg in pentaminomycins, as previously suggested (Kaweewan et al, 2020; Hwang et al, 2020). The pathway also contains regulatory genes and other genes of unknown function (Table 1).

**Table 1.** Closest BLAST homolog for each ORF in *cpp* BGC

| ORF | Lenght (aa) | Closest BLAST match | Identity(%) / Similarity(%) |
| --- | --- | --- | --- |
|  |  | [Organism]<br>GenBank reference |  |
| *cppA* | 502 | Hypothetical protein DEH18_05445<br>[*Streptomyces* sp. NHF165]<br>QHF93414.1 | 99/99 |
| *cppB* | 6187 | Non-ribosomal peptide synthase/polyketide synthase<br>[*Streptomyces cacaoi*]<br>WP_158102276.1 | 99/99 |
| *cppC* | 172 | MULTISPECIES: DUF2975 domain-containing protein<br>[*Streptomyces*]<br>WP_030891799.1 | 100/100 |
| *cppD* | 102 | Helix-turn-helix domain-containing protein<br>[*Streptomyces cacaoi*]<br>WP_149564434.1 | 99/99 |
| *cppE* | 420 | MULTISPECIES: sensor histidine kinase<br>[*Streptomyces*]<br>WP_051857187.1 | 99/100 |
| *cppF* | 280 | Hypothetical protein SCA03_67000<br>[*Streptomyces cacaoi* subsp. *cacaoi*]<br>GEB54149.1 | 100/100 |
| *cppG* | 131 | MULTISPECIES: DUF742 domain-containing protein<br>[*Streptomyces*]<br>WP_030891786.1 | 100/100 |
| *cppH* | 186 | MULTISPECIES: ATP/GTP-binding protein<br>[*Streptomyces*]<br>WP_030891784.1 | 100/100 |
| *cppI* | 451 | MULTISPECIES: cytochrome P450<br>[*Streptomyces*]<br>WP_030891781.1 | 100/100 |
| *cppJ* | 431 | Cytochrome P450<br>[*Streptomyces cacaoi*]<br>WP_086815207.1 | 99/99 |
| *cppK* | 271 | 3-hydroxybutyryl-CoA dehydratase<br>[*Streptomyces cacaoi* subsp. *cacaoi*]<br>GEB54144.1 | 99/99 |
| *cppL* | 141 | MULTISPECIES: hypothetical protein<br>[*Streptomyces*]<br>WP_141275837.1 | 100/100 |
| *cppM* | 5901 | Non-ribosomal peptide synthase/polyketide synthase<br>[*Streptomyces* sp. NHF165]<br>WP_159784853.1 | 99/99 |
| *cppN* | 69 | MULTISPECIES: hypothetical protein<br>[*Streptomyces*]<br>WP_030890086.1 | 100/100 |
| *cppO* | 175 | MULTISPECIES: hypothetical protein<br>[unclassified *Streptomyces*]<br>WP_030890089.1 | 99/100 |

**Figure 3.**
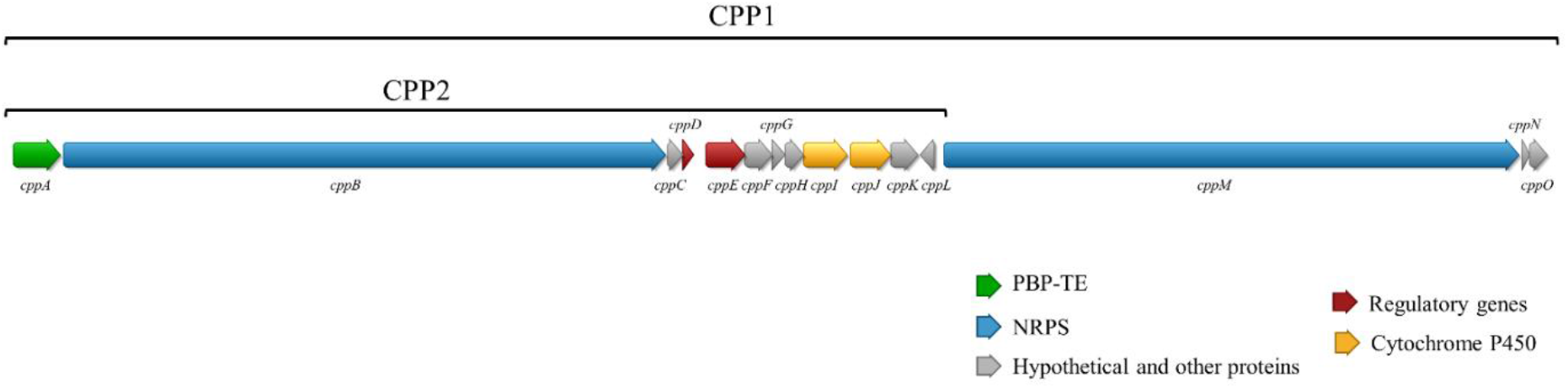
*cpp* biosynthetic gene cluster. It contains two NRPS genes (blue), one PBP-TE gene (green), two cytochromes P450 (yellow), two regulatory genes (red) and other genes with unknown functions (grey). The two fragments cloned by CATCH into vector pCAP01 are indicated: the 28.7 Kb CPP2 fragment contains the PBP-like protein, the NRPS1(*cppB*) and the genes present between NRPS1and NRPS2; the 48 Kb CPP1 fragment includes the above described 28.7 Kb fragment and the NRPS2 (*cppM*) gene.

**Figure 4.**
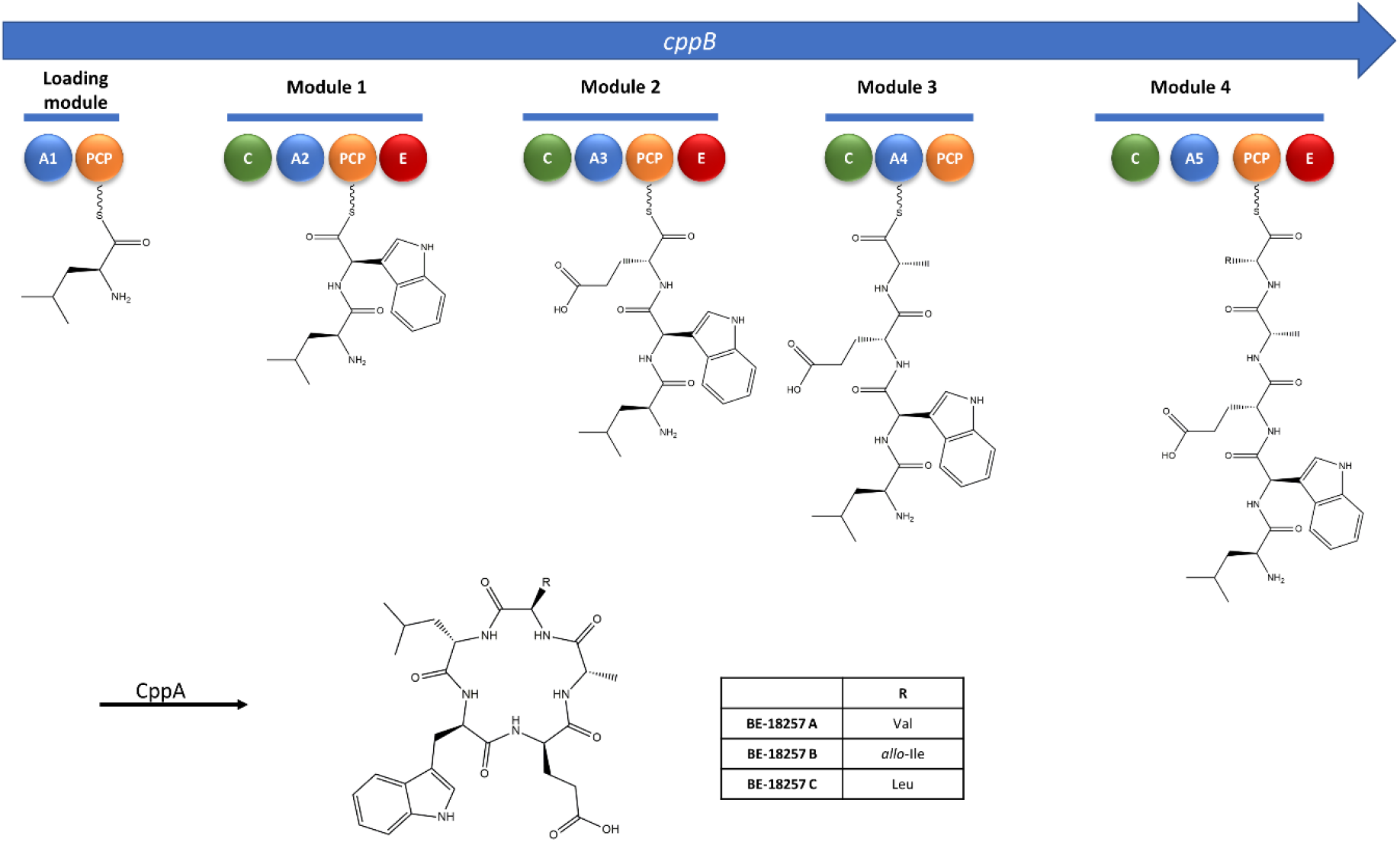
Proposed biosynthetic pathway for the BE-18257 A-C antibiotics with the nonribosomal peptide synthetase CppB modular organization. A, adenylation domain; PCP, peptidyl carrier protein; C, condensation domain; E, epimerase domain; CppA, PBP-TE.

**Figure 5.**
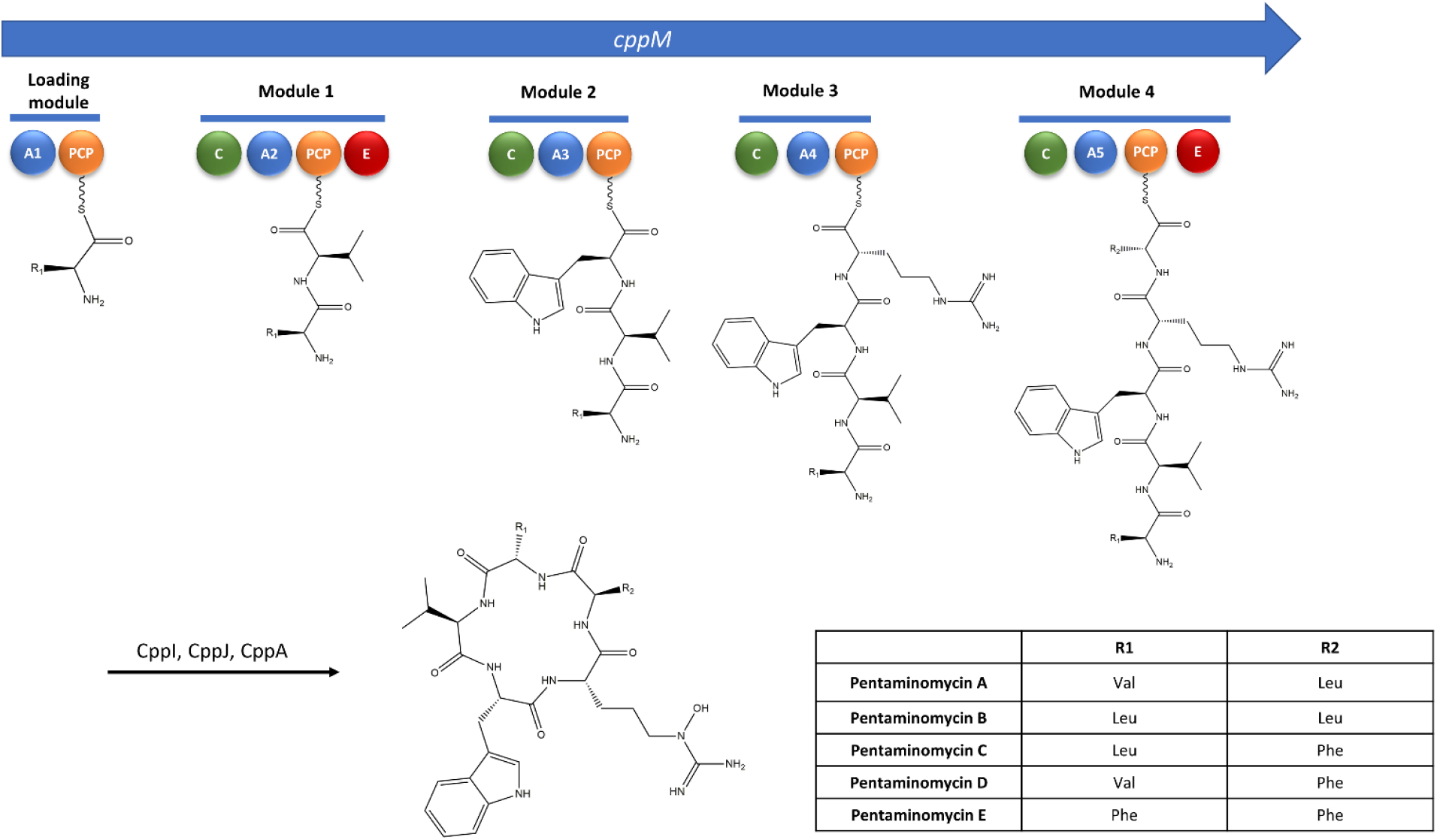
Proposed biosynthetic pathway for the pentaminomycins A-E with the nonribosomal peptide synthetase CppM modular organization. A, adenylation domain; PCP, peptidyl carrier protein; C, condensation domain; E, epimerase domain; CppI and CppJ, cytochromes P450; CppA, PBP-TE.

### 4.3. Cloning and heterologous expression of the *cpp* gene cluster

To demonstrate that the identified *cpp* cluster is involved in the biosynthesis of both BE-18257 A-C and pentaminomycins A-E, we separately cloned two different fragments of the BGC by Cas9-assisted targeting of chromosome segments (CATCH) cloning (Jiang et al, 2015), a main approach to clone long microbial genomic sequences, into vector pCAP01 (Yamanaka et al, 2014). This method uses in-gel RNA-guided Cas9 nuclease digestion of bacterial DNA, which is subsequently ligated with cloning vector by Gibson assembly (Jiang & Zhu, 2016). The first genome sequence cloned was a 28.7 Kb fragment containing the PBP-like protein gene (*cppA*), the NRPS1 gene (*cppB*) and the genes present between NRPS1 and NRPS2 (*cppC-L*) to obtain pCPP1; the second one was a 48 Kb fragment including all the genes supposed to be required for the biosynthesis of both antibiotics; this was the previously described 28.7 Kb fragment and the NRPS2 gene (*cppM*), together with two genes enconding hypothetical proteins downstream NRPS2 (*cppN-O*), to obtain pCCP2 (Figure 3).

The plasmids pCPP1 and pCPP2, were transformed into *E. coli* NEB 10-beta competent cells. Clones were checked by restriction analysis, and one of the clones harboring pCPP1 and another one harboring pCPP2 were selected to perform intergeneric conjugations. Since pCPP1 and pCPP2 contain the kanamycin-resistant marker, we could not directly transform non-methylating Cm^R^ Km^R^ *E. coli* ET12567/pUB307. Thus, we performed two triparental intergeneric conjugations using *E. coli* NEB-10/pCPP1 and ET12567/pUB307 or *E. coli* NEB-10/pCPP2 and ET12567/pUB307 as donor strains, and spores of *S. albus* J1074 as recipient strain. For the negative control, a triparental conjugation was also made using *E. coli* NEB-10/pCAP01 and ET12567/pUB307 as donor strains and the same recipient strain. Transconjugants were checked by PCR with primers BLAC check-F and BLAC check-R (Supporting Table 1) to confirm the integration of the cloned BGCs into the chromosome of *S. albus* J1074.

Five positive transconjugants from each conjugation, together with the negative control and the wild-type strain *S. cacaoi* CA-170360, were grown in liquid MPG and R2YE media (to favor the detection of BE-18257 antibiotics and pentaminomycins, respectively) during 14 days at 28°C, and acetone extracts from the cultures whole broths were prepared. After removing the solvent, the residue was resuspended in 20% DMSO/water and analyzed by LC-HRESI-TOF.

The analysis of extracts from pCPP1 and pCPP2 transconjugants confirmed the presence of peaks at 3.46 minutes and 3.77 minutes, coincident with the retention time of elution of the three BE-18257 A-C isolated from the CA-170360 strain (Supporting Figures 2 and 3). The detection levels of the BE-18257 A-C molecules in the pCPP1 transconjugants (which lacked the pentaminomycins NRPS gene) were much higher than in the pCPP2 transconjugants (which carried in addition the pentaminomycins NRPS gene). The analysis of the pCPP2 transconjugant also confirmed the presence of peaks coincident with the retention times of elution of the pentaminomycins D and E, isolated from CA-170360, which were absent in the pCPP1 transconjugants (Supporting Figures 4 and 5).

The correlation between the UV spectrum, exact mass and isotopic distribution between the BE-18257 and pentaminomycins from *S. cacaoi* CA-170360 (Supporting Figures 2-6) and the components isolated from the transconjugants *S. albus*/pCPP1 and *S. albus*/pCPP2 (Supporting Figures 2-5) unequivocally confirmed that they corresponded to BE-18257 A-C in the case of *S. albus*/pCPP1 and to BE-18257 A-C and pentaminomycins D and E in the case of *S. albus*/pCPP2. In the pCPP2 transconjugants, we detected ions suggesting the presence of pentaminomycins A, B and C, but given the low production levels of these compounds (data not shown) we could not obtain proper mass spectra.The detection levels of all the cyclopentapeptides in the heterologous hosts is lower than in the *S. cacaoi* strain, in which the pentaminomycins A-E were already poorly produced. Consequently, the productions of pentaminomycins in the heterologous host *S. albus*/pCPP2 was still at the limit of detection from most of the compounds. These results clearly demonstrate that the first NRPS gene (*cppB*) is the responsible of the biosynthesis of BE-18257 antibiotics, and that the second NRPS gene (*cppM*) synthesizes pentaminomycins. The results also suggest that the cluster-located PBP-TE is involved in the cyclization of both compounds, and that the *cpp* BGC can be considered an atypical case in which two types of independent compounds are processed by the same enzyme.

The genome of *S. cacaoi* CA-170360 also contains some genes related to tryptophan biosynthesis downstream from the second NRPS gene (*cppM*) (data not shown), as it has been already described by Hwang et al (2020) for strain GG23. As both pentaminomycins and BE-18257 contain tryptophan in their structures, it has been proposed that they may share the Trp biosynthetic genes to incorporate this amino acid (Hwang et al, 2020). However, our results clearly show that those genes are not required to incorporate Trp in the cyclic pentapeptides, since they were not included in the fragment cloned into pCPP1 and in pCPP2, and the pentaminomycins and BE-18257 were still produced. This indicates that the tryptophan, as well as the rest of amino acids, are obtained from the primary metabolism amino acid pool.

### 4.4. Genome mining of *cpp*-like BGCs

A tblastn search of the NRPS1 and NRPS2 protein sequences against both nucleotide and Whole Genome Sequence (WGS) databases from NCBI showed that the *cpp* cluster is also present in some genomes described in Figure 6. Moreover, the pathway is only present in strains belonging to *Streptomyces cacaoi* species: *S. cacaoi* NHF-165, *S. cacaoi* DSM40057, *S. cacaoi* subs. *cacaoi* NRRL-1220, *S. cacaoi* OABC16, *S. cacaoi* NRRL S-1868, *S. cacaoi* NRRL F-5053 and *S. cacaoi* NBRC 12748 (Figure 6). The *pen* cluster described by Hwang et al (2020) in the strain *Streptomyces* sp. GG23, which has been also identified as a strain of *Streptomyces cacaoi,* has not been included in this analysis because the sequence is not yet available. Nevertheless, the comparison of the homologies described in the *pen* and in the *cpp* clusters clearly shows that they are highly similar. This indicates, as well as it was described for the cacaoidin cluster (Román-Hurtado et al, 2020), that the *cpp* cluster is highly conserved within members of this species and is another excellent example of the biosynthesis of a specialized metabolite that could be used as a species-specific trait (Vicente et al, 2018).

**Figure 6.**
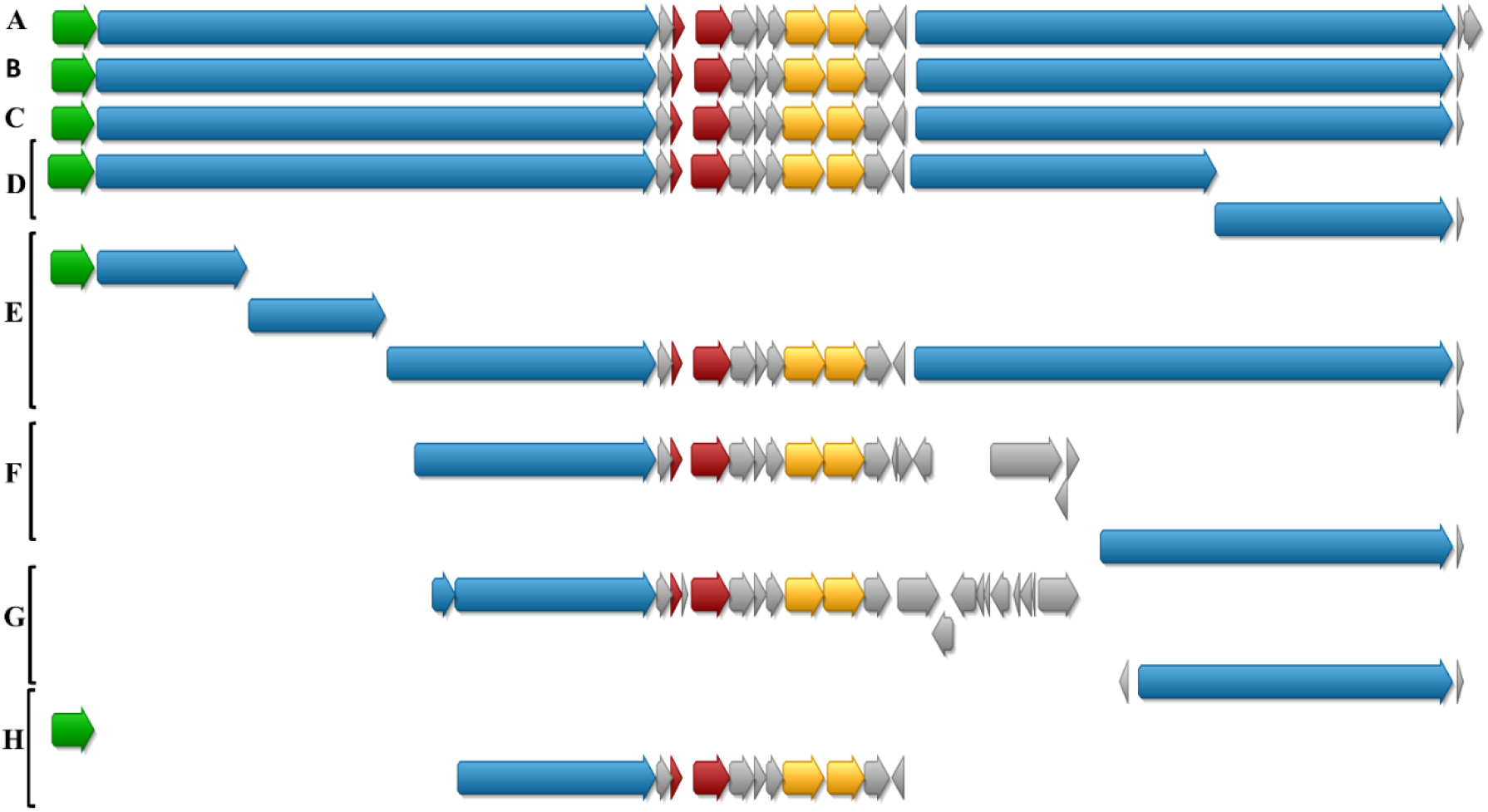
Schematic representation of the alignment of *cpp* BGC from *Streptomyces cacaoi* CA-170360 and the homologous genome sequences found in NCBI. A, *S. cacaoi* CA-170360; B*, S. cacaoi* NHF165; C, *S. cacaoi* DSM40057; D, *S. cacaoi* NRRL B-1220; E, *S. cacaoi* OABC16; F, *S. cacaoi* NRRL S-1868; G, *Streptomyces* sp. NRRL F-5053 and H, *S. cacaoi* NBRC 12748. Due to the high fragmentation in some of these *S. cacaoi* genomes, the corresponding BGC was found in different contigs and was not complete.

## 5. Conclusions

We have shown that pentaminomycins and BE-18257 antibiotics can be produced heterologously from a single BGC (*cpp*) containing two independent NRPS genes, *cppB* and <u>*cppM*</u>, encoding respectively each type of pentapeptide, and one PBP-like stand-alone protein (CppA) that is proposed to be involved in the release and cyclization of both families of compounds. We have also demonstrated that the downstream genes related to tryptophan biosynthesis, an amino acid that is present in all the cyclic pentapeptides, are not necessary for their production. Furthermore, our bioinformatic analysis suggest that the *cpp* cluster might be a species-specific trait since was only found in the genomes of all publicly available *Streptomyces cacaoi* strains and not in other species. Despite the lack of similar BGCs found in genome sequence databases beyond *Streptomyces cacaoi* species, this work opens the door to identify additional tandem biosynthetic gene organized within a single BGC to ensure the biosynthesis or related families of compounds, as well as the biosynthesis of new analogs of both BE-18257 antibiotics and pentaminomycins.

## 7. Acknowledgements

This work is supported by Novo Nordisk Foundation grant NNF16OC0021746. The authors thank Daniel Oves-Costales for helpful advice during the whole process and the Microbiology and Chemistry areas of Fundación MEDINA for the technical support. We thank Bradley Moore for providing plasmid pCAP01. We thank José Antonio Salas and the University of Oviedo for kindly providing strains *Streptomyces albus* J1074 and *Escherichia coli* ET12567/pUB307.

## 8. Author Contributions

F.R.H, M.S.H. and O.G. conceived and designed the experiments and wrote the paper. F.R.H. performed the genetic experiments and bioinformatic analysis. J.M. performed the LC-HRESI-TOF experiments. F.J.O.L., D.C.M. and F.R. confirmed the structures of the compounds . All authors reviewed and approved the manuscript.

