## Supplementary Information for "One pathway, two cyclic pentapeptides: heterologous expression of BE-18257 A-C and pentaminomycins A-E from *Streptomyces cacaoi* CA-170360"

Fundación MEDINA, Avenida del Conocimiento 34, 18016 Granada, Spain

*Corresponding autor:

### SUPPORTING ON-LINE INFORMATION

#### Culture media composition

#### ATCC-2

- Soluble starch 20 g/L
- Glucose 10 g/L
- NZ Amine Type E 5 g/L
- Meat extract 3 g/L
- Peptone 5 g/L
- Yeast extract 5 g/L
- Calcium carbonate 1 g/L

  pH adjusted to 7

#### YEME

- Yeast extract 3 g/L
- Bacto-peptone 5 g/L
- Oxoid malt extract 3 g/L
- Glucose 10 g/L
- Sucrose 340 g
- $MgCl_2$-$6H_2O$ 2mL/L

**R2YE**

- Yeast extract 5 g/L
- Sucrose 103 g/L
- $K_2SO_4$ 0.25 g/L
- $MgCl_2$-6$H_2O$ 10.12 g/L
- Glucose 10 g/L
- Casamino acids 0.1 g/L
- $KH_2PO_4$ 0.5% 1 mL
- $CaCl_2$-2$H_2O$ 3.68% 8 mL
- L-proline 20% 1.5 mL
- TES buffer 5.73% adjusted to pH7.2 10 mL
- Trace element solution 0.2 mL
- NaOH 1N 0.5mL
- Growth factors for auxotrophs 0.75 mL

Trace element solution:

- $ZnCl_2$ 40 mg/L
- $FeCl_3$-6$H_2O$ 200 mg/L
- $CuCl_2$-2$H_2O$ 10 mg/L
- $MnCl_2$-4$H_2O$ 10 mg/L
- $Na_2B_4O_7$-10$H_2O$ 10 mg/L
- $(NH_4)_6Mo_7O_{24}$-4$H_2O$ 10 mg/L

**KM4**

- Glucose 4 g/L
- Yeast extract 4 g/L
- Malt extract 10 g/L
- $CaCO_3$ 2 g/L

**MPG**

- Glucose 10 g/L
- Millet 20 g/L
- Cottonseed flour 20 g/L
- MOPS 20g/L

  pH adjusted to 7

**FR23**

- Glucose 5 g/L
- Soluble starch from potato 30 g/L
- Cottonseed flour 20 g/L
- Cane molasses 20 g/L

  pH adjusted to 7

**DEF-15**

- Sucrose 40 g/L
- $ClNH_4$ 2 g/L, $Na_2SO_4$ 2 g/L
- $K_2HPO_4$ 1 g/L
- $Cl_2Mg.6H_2O$ 1 g/L
- Trace elements 1mL
- $CaCO_3$ 2 g/L

  pH adjusted to 7

Trace elements:

- $MnCl_2$-$4H_2O$ 0.1/100 mL
- $ZnCl_2$ 0.1 g/100 mL
- $FeCl_2$-$4H_2O$ 0.1 g/100 mL
- NaI 0.05 g/100 mL

**MA**

- MOPS 21 g/L
- Glucose 5 g/L
- Yeast extract 0.5 g/L
- Beef extract 0.5 g/L
- Casamino acids 1 g/L
- Agar 25 g/L

pH adjusted to 7

### <u>Figures</u>

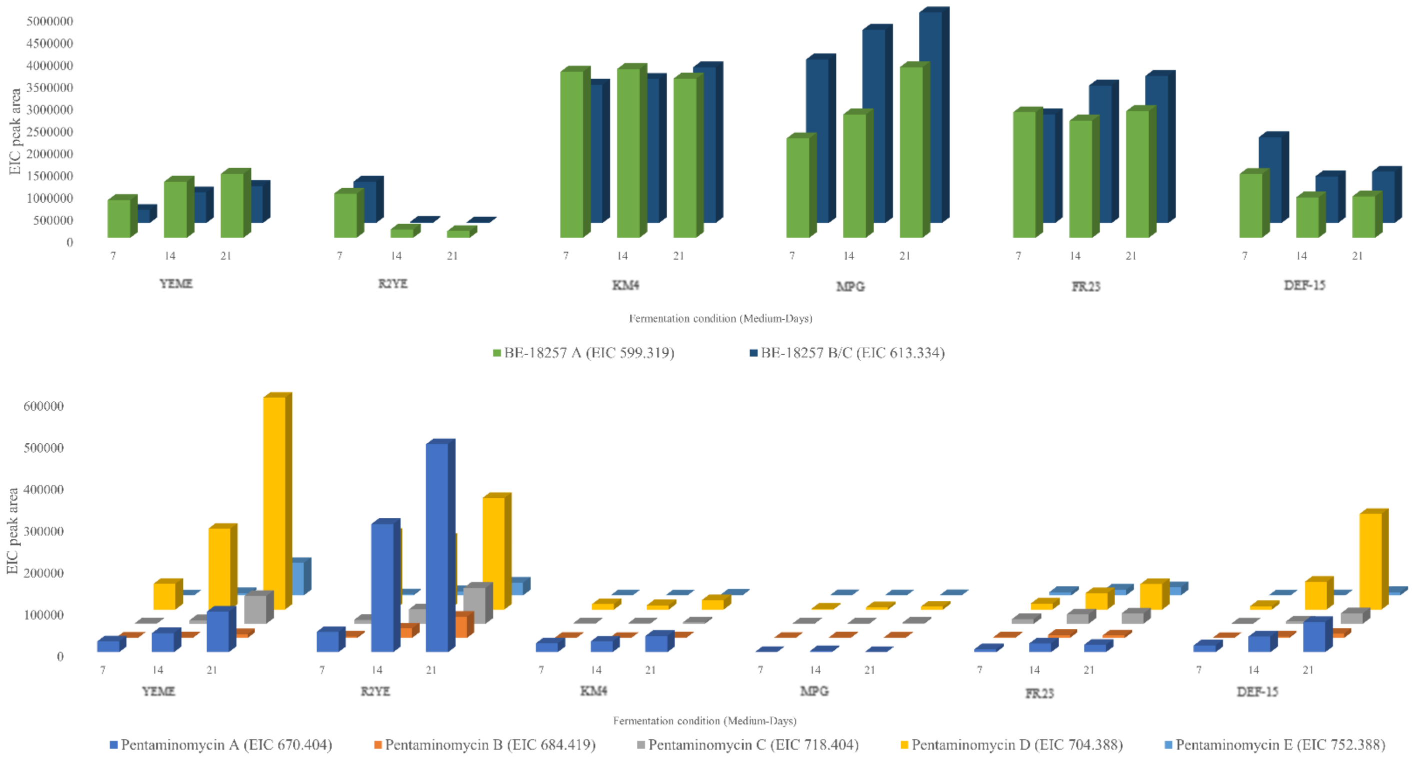


**Supporting Figure 1.** Production of BE-18257 antibiotics (top) and pentaminomycins A-E (bottom) by strain *S. cacaoi* CA-170360 in six different media at three different times (7, 14 and 21 days). The average extracted ion chromatogram (EIC) peak area from triplicate culture extracts of the strain CA-170360 is represented.

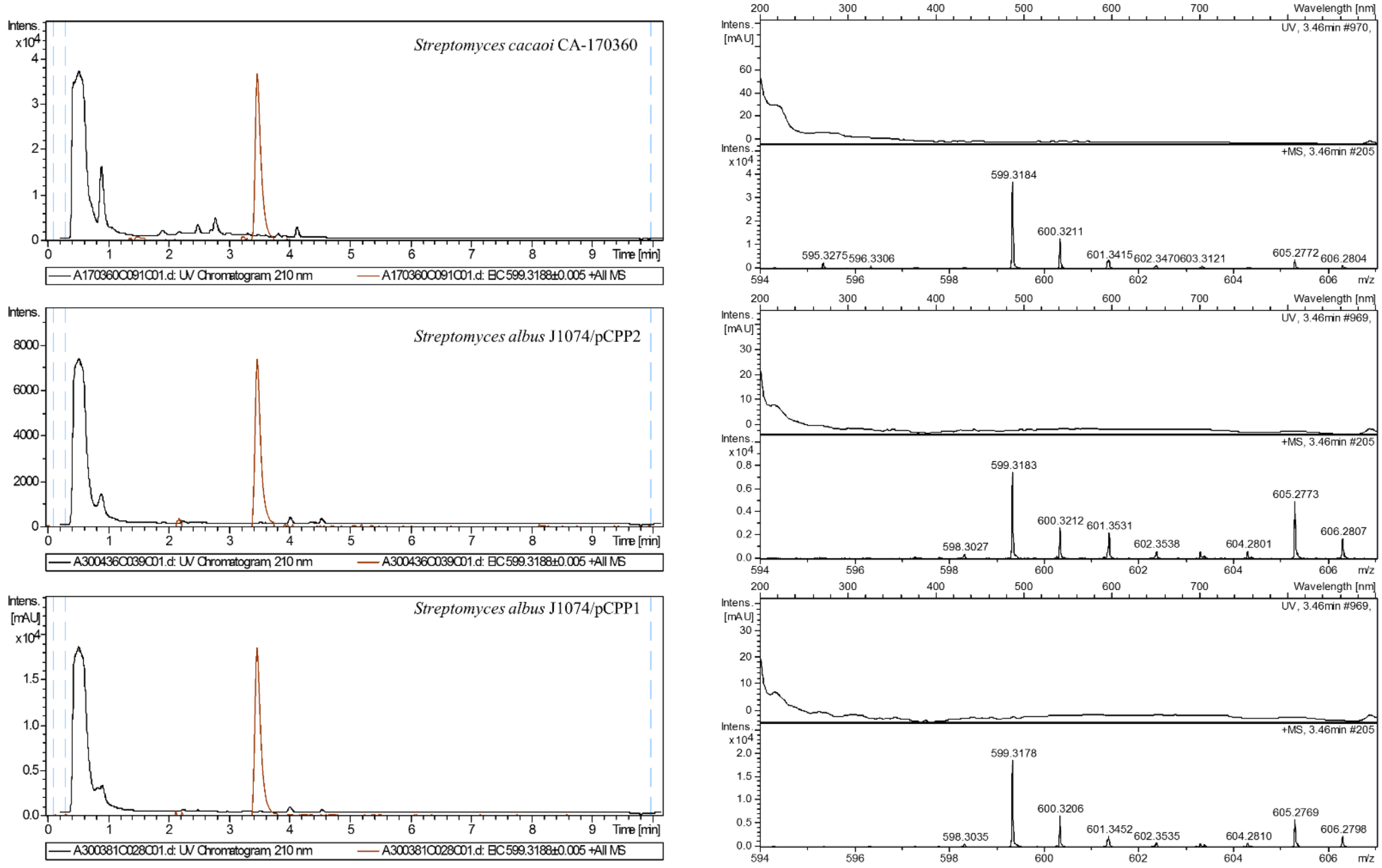


**Supporting Figure 2.** BE-18257 A production. Left. Chromatograms of UV absorbance at 210 nm and extracted ion m/z = 599.3188 ± 0.005, $C_{30}H_{43}N_6O_7^+$ of BE-18257 A from original producing strain *Streptomyces cacaoi* CA-170360 (top) and the heterologous producing strains *Streptomyces albus* J1074/pCPP2 (middle) and *Streptomyces albus* J1074/pCPP1 (bottom). Right. Experimental UV and positive mass spectra from $C_{30}H_{43}N_6O_7^+$ (calculated value: 599.3188) adduct from original producing strain *Streptomyces cacaoi* CA-170360 (top) and the heterologous producing strains *Streptomyces albus* J1074/pCPP2 (middle) and *Streptomyces albus* J1074/pCPP1 (bottom).

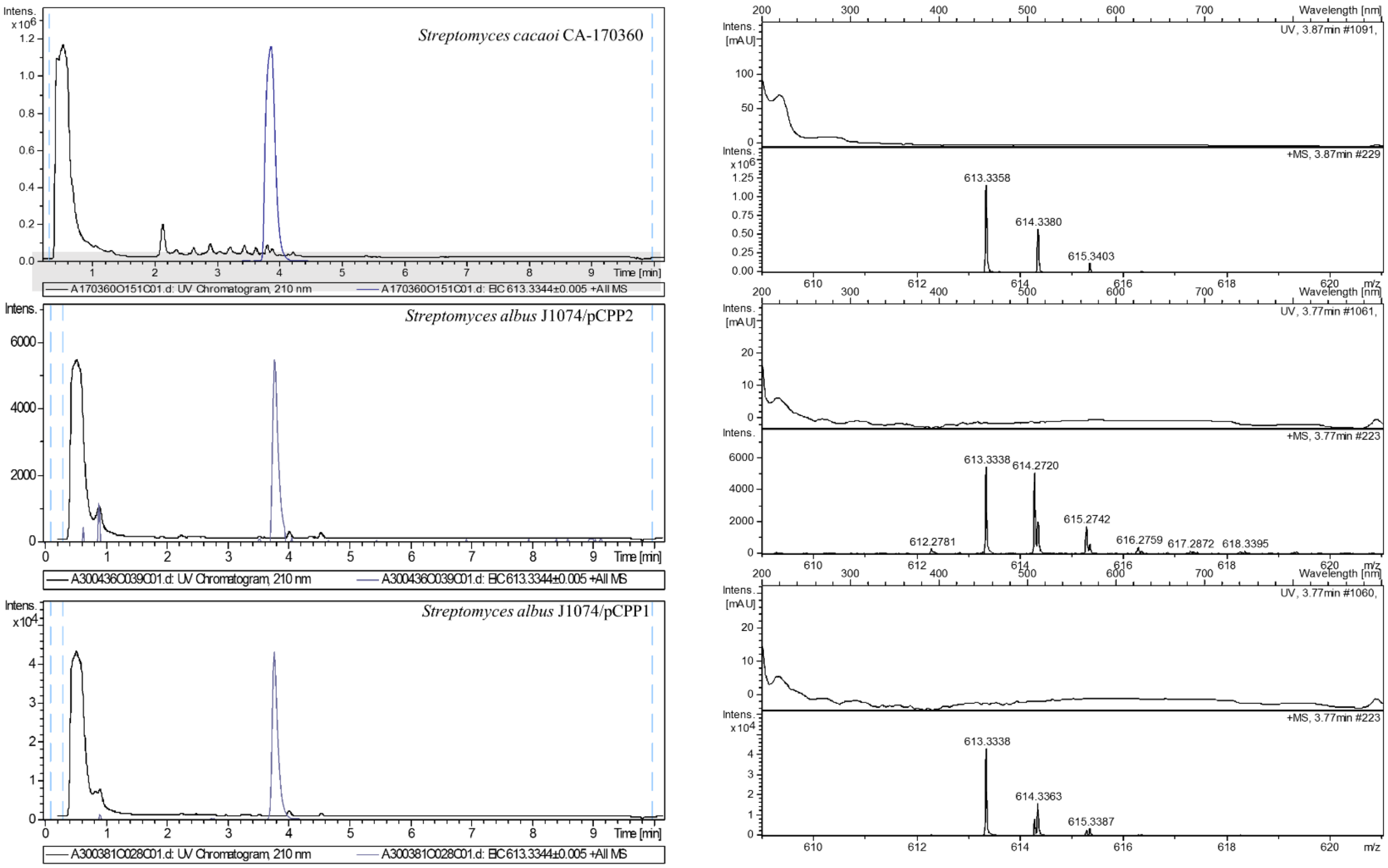


**Supporting Figure 3.** BE-18257 B/C production. Left. Chromatograms of UV absorbance at 210 nm and extracted ion m/z = 613.3344 ± 0.005, $C_{31}H_{45}N_6O_7^+$ of BE-18257 B/C from original producing strain *Streptomyces cacaoi* CA-170360 (top) and the heterologous producing strains *Streptomyces albus* J1074/pCPP2 (middle) and *Streptomyces albus* J1074/pCPP1 (bottom). Right. Experimental UV and positive mass spectra from $C_{31}H_{45}N_6O_7^+$ (calculated value: 613.3344) adduct from original producing strain *Streptomyces cacaoi* CA-170360 (top) and the heterologous producing strains *Streptomyces albus* J1074/pCPP2 (middle) and *Streptomyces albus* J1074/pCPP1 (bottom).

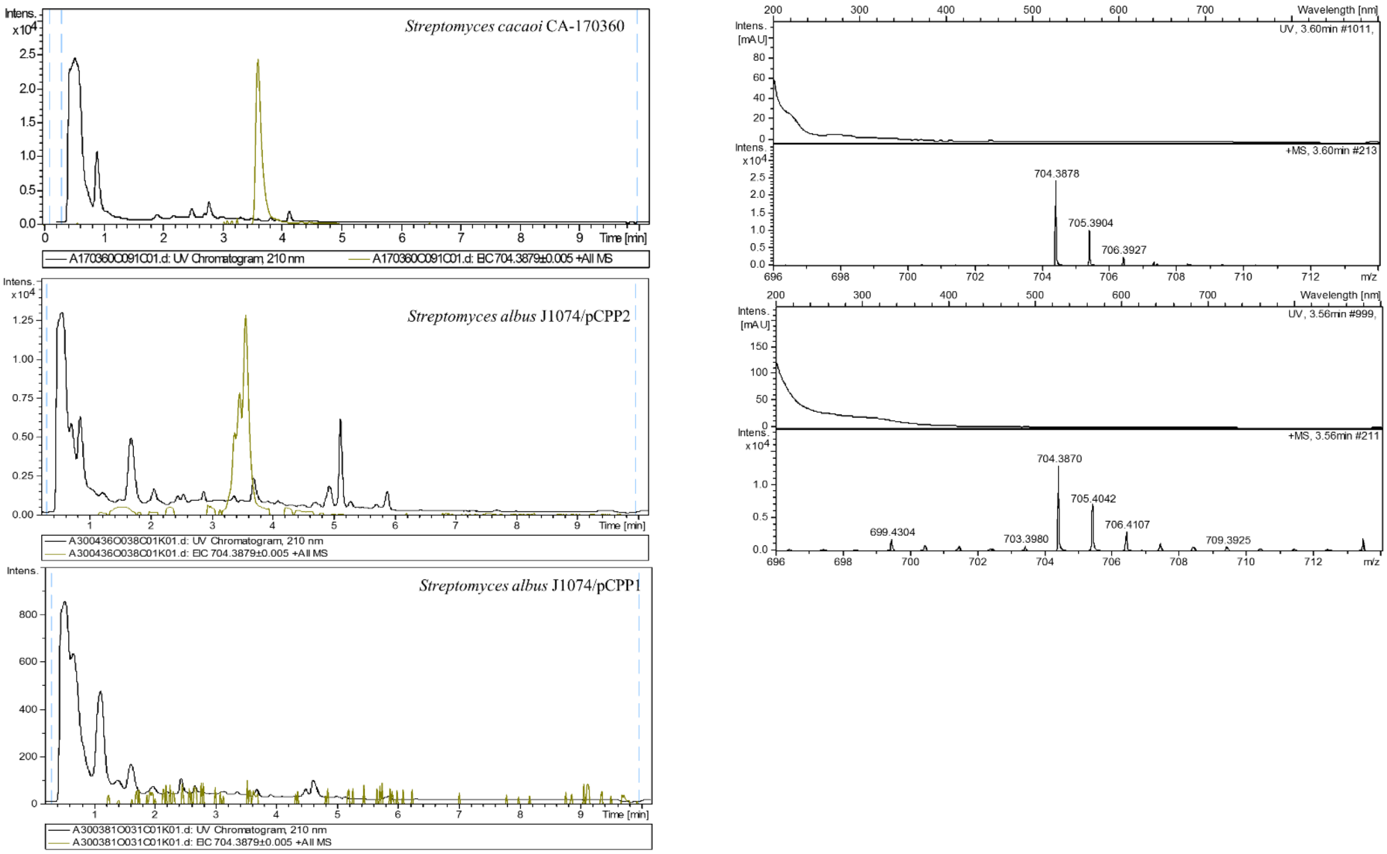


**Supporting Figure 4.** Pentaminomycin D production. Left. Chromatograms of UV absorbance at 210 nm and extracted ion m/z = 704.3879 ± 0.005, $C_{36}H_{50}N_9O_6^+$ of pentaminomycin D from original producing strain *Streptomyces cacaoi* CA-170360 (top) and the heterologous producing strains *Streptomyces albus* J1074/pCPP2 (middle) and *Streptomyces albus* J1074/pCPP1 (bottom). Right. Experimental UV and positive mass spectra from $C_{36}H_{50}N_9O_6^+$ (calculated value: 704.3879) adduct from original producing strain *Streptomyces cacaoi* CA-170360 (top) and the heterologous producing strain *Streptomyces albus* J1074/pCPP2 (middle). No UV or mass spectra was obtained with the heterologous producing strain *Streptomyces albus* J1074/pCPP1 as it did not carry the NRPS gene required for the production of pentaminomycins.

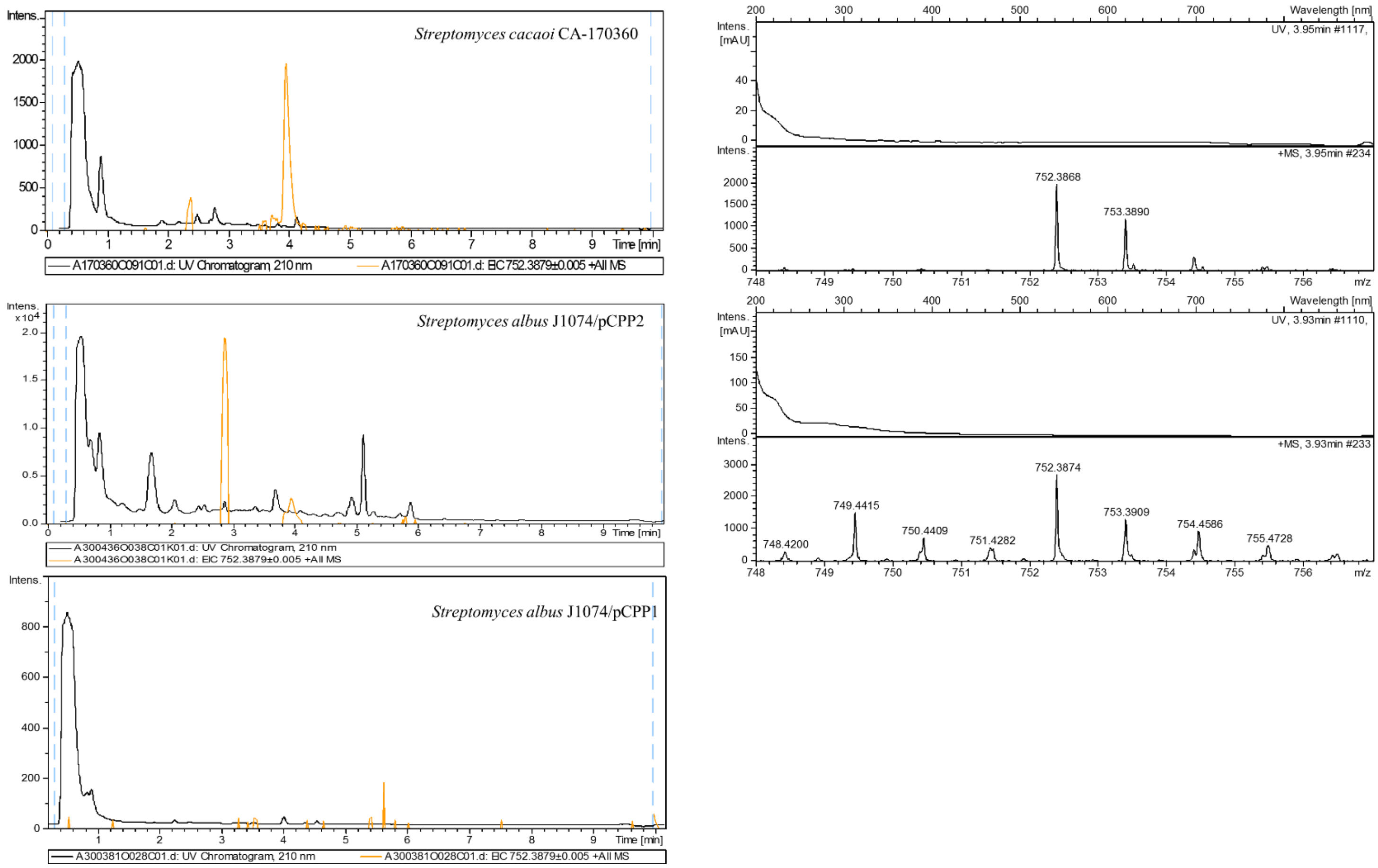


**Supporting Figure 5.** Pentaminomycin E production. Left. Chromatograms of UV absorbance at 210 nm and extracted ion m/z = 752.3879 ± 0.005, $C_{40}H_{50}N_9O_6^+$ of pentaminomycin E from original producing strain *Streptomyces cacaoi* CA-170360 (top) and the heterologous producing strains *Streptomyces albus* J1074/pCPP2 (middle) and *Streptomyces albus* J1074/pCPP1 (bottom). Right. Experimental UV and positive mass spectra from $C_{40}H_{50}N_9O_6^+$ (calculated value: 752.3879) adduct from original producing strain *Streptomyces cacaoi* CA-170360 (top) and the heterologous producing strain *Streptomyces albus* J1074/pCPP2 (middle). No UV or mass spectra was obtained with the heterologous producing strain *Streptomyces albus* J1074/pCPP1 as it did not carry the NRPS gene required for the production of pentaminomycins.

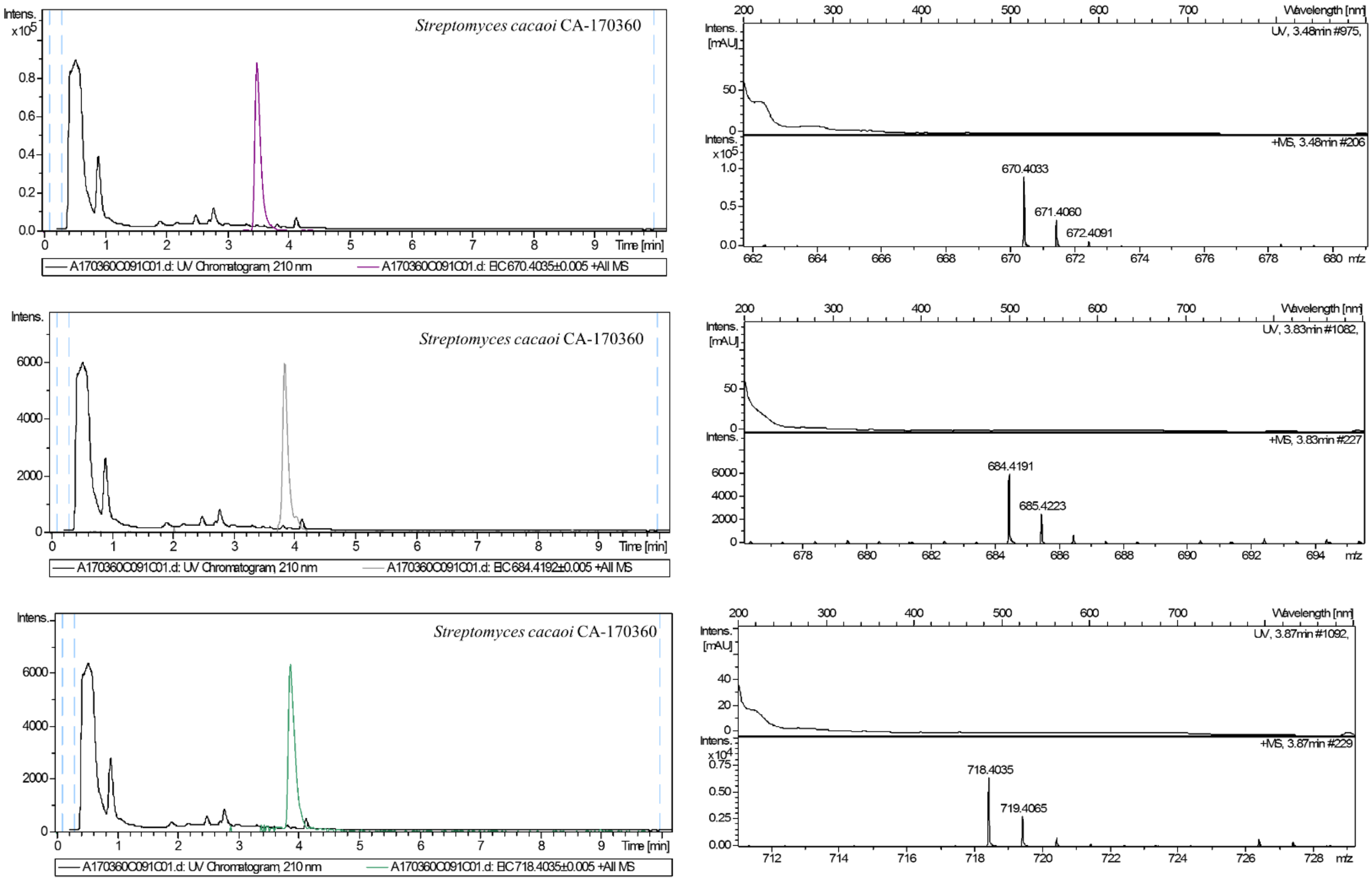


**Supporting Figure 6.** Pentaminomycins A, B and C production. Left. Chromatogram of UV absorbance at 210 nm and extracted ion m/z = 670.4035 ± 0.005, $C_{33}H_{52}N_9O_6^+$ of pentaminomycin A (top), extracted ion m/z = 684.4192 ± 0.005, $C_{34}H_{54}N_9O_6^+$ of pentaminomycin B (middle) and extracted ion m/z = 718.4035 ± 0.005, $C_{37}H_{52}N_9O_6^+$ of pentaminomycin C (bottom) from original producing strain *Streptomyces cacaoi* CA-170360. Right. Experimental UV and positive mass spectra from $C_{33}H_{52}N_9O_6^+$ (calculated value: 670.4035) adduct (top), $C_{34}H_{54}N_9O_6^+$ (calculated value: 684.4192) adduct (middle) and $C_{37}H_{52}N_9O_6^+$ (calculated value: 718.4035) adduct (bottom) from original producing strain *Streptomyces cacaoi* CA-170360. Proper mass or ultraviolet spectra of pentaminomycins A, B and C could not be obtained in the heterologous hosts.

### Tables

| **Oligonucleotide** | **Sequence (5’-3’)** |
|---|---|
| **Penta1-sgRNA** | TAATACGACTCACTATAGATGATCCAGAATCCGTGCTTGTTTTAGAGCTAGAAATAGCAA |
| **Penta2-sgRNA** | TAATACGACTCACTATAGACCCCAGACTTCAGCGTTTGGTTTTAGAGCTAGAAATAGCAA |
| **Penta3-sgRNA** | TAATACGACTCACTATAGGAACTGAAGGCACAACCAAAGTTTTAGAGCTAGAAATAGCAA |
| **sgRNA-F** | GTTTTAGAGCTAGAAATAGCAAGTTAAAATAAGGCTAGTC |
| **sgRNA-R** | AAAAGCACCGACTCGGTGCCACTTTTTCAAGTTGATAACGGACTAGCCTTATTTTAACT |
| **pCAP01-Penta1-F** | AGGCTAGTCAGGGGTACCGGGCCCCTCAAATCGAGACTTGAGGTACCTGT |
| **pCAP01-Penta1-R** | TCGGAAAGCGGCTGAAGGTCTCTCCCAAGCTCGAGGTTACTAGTCGATCT |
| **pCAP01-Penta2-F** | AAGGCTAGTCAGGGGTACCGGGCCCCTCAATCGAGACTTGAGGTACCTGT |
| **pCAP01-Penta2-R** | GGCCAACTGGCCTGCTACCTGCGCCATTTGTCGAGGTTACTAGTCGATCT |
| **BLAC check-F** | CCAACTCCTCGAACAGCT |
| **BLAC check-R** | CTGCTCAGCCACACCG |

**Supporting Table 1.** Oligonucleotides used in this work
